# Laser-driven VHEE pulsed fast fractionation (PFF): second-scale inter-pulse timing can modulate normal tissue toxicity in vivo

**DOI:** 10.64898/2025.12.24.696225

**Authors:** Camilla Giaccaglia, Emilie Bayart, Chaitanya Varma, Jean-Philippe Goddet, Julien Gautier, Amar Tafzi, Emilie Menant, Isabelle Lamarre, Sophie Heinrich, Charles Fouillade, Alessandro Flacco

## Abstract

Radiotherapy (RT) is constrained by the narrow therapeutic window between tumour control and normal-tissue toxicity. While FLASH RT has been reported to reduce normal-tissue toxicity at ultra-high dose rates (UHDR) while preserving tumour control in several preclinical models, the radiobiological potential of intermediate temporal regimes between conventional fractionation and FLASH remains largely unexplored. Here, we show that temporal separation of ultrashort (fs–ps) dose-delivery pulses delivered at ultra-high instantaneous dose rate, with second-scale inter-pulse intervals, can differentially modulate biological response in vitro and in vivo. Samples were irradiated with very high-energy electrons (VHEEs, 50 MeV to 300 MeV) generated by a laser-plasma accelerator (LPA), using a temporal irradiation modality we term “pulsed fast fractionation” (PFF), with instantaneous dose rates exceeding 10^12^ Gy s^−1^.

Varying the inter-pulse interval from 1 s to 10 s while maintaining dose per pulse and total dose constant modulated biological response in both in vitro and in vivo models. At the shortest investigated inter-pulse interval (*θ* = 1 s), MRC5-hTERT human fibroblasts showed higher viability, whereas HCT116 colorectal carcinoma cells showed lower viability under matched dose conditions. In vivo, the same interval was associated with reduced radiation-induced growth impairment in zebrafish embryos, consistent with the response pattern observed in the non-tumour cell model and extending this timing-dependent response to a whole-organism context. These findings identify second-scale pulse timing as a biologically active degree of freedom for VHEE delivery, positioning PFF as a distinct temporal irradiation regime with the potential to complement spatial dose modulation through temporal optimisation and extend the current fractionation-FLASH framework.

## I. INTRODUCTION

Radiotherapy (RT) is a cornerstone of cancer treatment, with over half of all patients receiving it during the course of their care [1]. Its therapeutic efficacy relies on achieving tumour control while limiting toxicity to surrounding normal tissues. Advances in treatment planning and beam delivery have progressively improved this balance through increasingly precise spatial control of dose, while temporal optimisation has traditionally been implemented at the macroscopic scale through fractionation. Conventional fractionation typically distributes the prescribed dose over daily fractions delivered across several weeks, exploiting differences in repair, re-oxygenation, redistribution, and repopulation between normal and tumour tissues [2, 3]. Alternative fractionation schemes modify dose per fraction and treatment schedule, but remain defined on macroscopic timescales of hours to days [4]. More recently, the observation of the FLASH effect [5] has shown that biological response can also differ when radiation is delivered over much shorter timescales, typically within hundreds of milliseconds and at ultra-high average dose rates on the order of 40 Gy s^*−*1^ or higher, with preferential normal-tissue sparing relative to tumour response. These observations have broadened the temporal framework of radiobiology beyond macroscopic fractionation, drawing attention to how dose is organised within an individual irradiation.

When considering the temporal organisation of dose within an irradiation, pulsed beam delivery can be characterised not only by total dose, average dose rate, and overall irradiation time, but also by the properties and spacing of individual pulses, including pulse duration (*τ*), dose per pulse (*D*_*p*_), and inter-pulse interval (*θ*). These parameters describe how the delivered dose is distributed across different temporal scales. In conventional clinical electron delivery, radiation is administered as a high-repetition pulse train with closely spaced pulses, such that dose deposition is effectively quasi-continuous on the second timescale. FLASH irradiation, although encompassing a range of pulse structures, is generally associated with ultra-high average dose rates and short overall irradiation times. Comparatively little attention has been given to a different combination of temporal characteristics: ultrashort dose-deposition events separated by second-scale inter-pulse intervals. Under such conditions, ultra-high instantaneous dose rates coexist with an irradiation extending over several seconds, while the average dose rate may remain below the range typically associated with FLASH.

Experimental investigation of this temporal regime is constrained by the availability of radiation sources capable of accessing these delivery conditions. Laser-plasma accelerators (LPAs) are particularly suited to this purpose, as they can generate femto-to picosecond electron bunches with instantaneous dose rates exceeding 10^9^ Gy s^*−*1^, while allowing controlled inter-pulse intervals on the second timescale. Their gigavolt-per-metre acceleration gradients further enable the production of very high-energy electrons (VHEEs, *E >* 50 MeV) over millimetre-scale distances. VHEEs offer deep penetration, favourable transport properties, and reduced sensitivity to tissue heterogeneities [6, 7], motivating growing interest in LPA-driven VHEE sources for radiotherapy research [8].

Evidence that second-scale timing of ultra-high instantanous dose-rate pulses can modulate biological response in vitro has previously emerged from our work with laser-driven protons [9]. We refer to this irradiation regime as “pulsed fast fractionation” (PFF): ultrashort pulses delivered at controlled second-scale inter-pulse intervals under ultra-high instantaneous dose-rate conditions. The relative positioning of PFF with respect to representative conventional and FLASH linac-based electron delivery is summarised schematically in Figure 1.

**FIG. 1.**
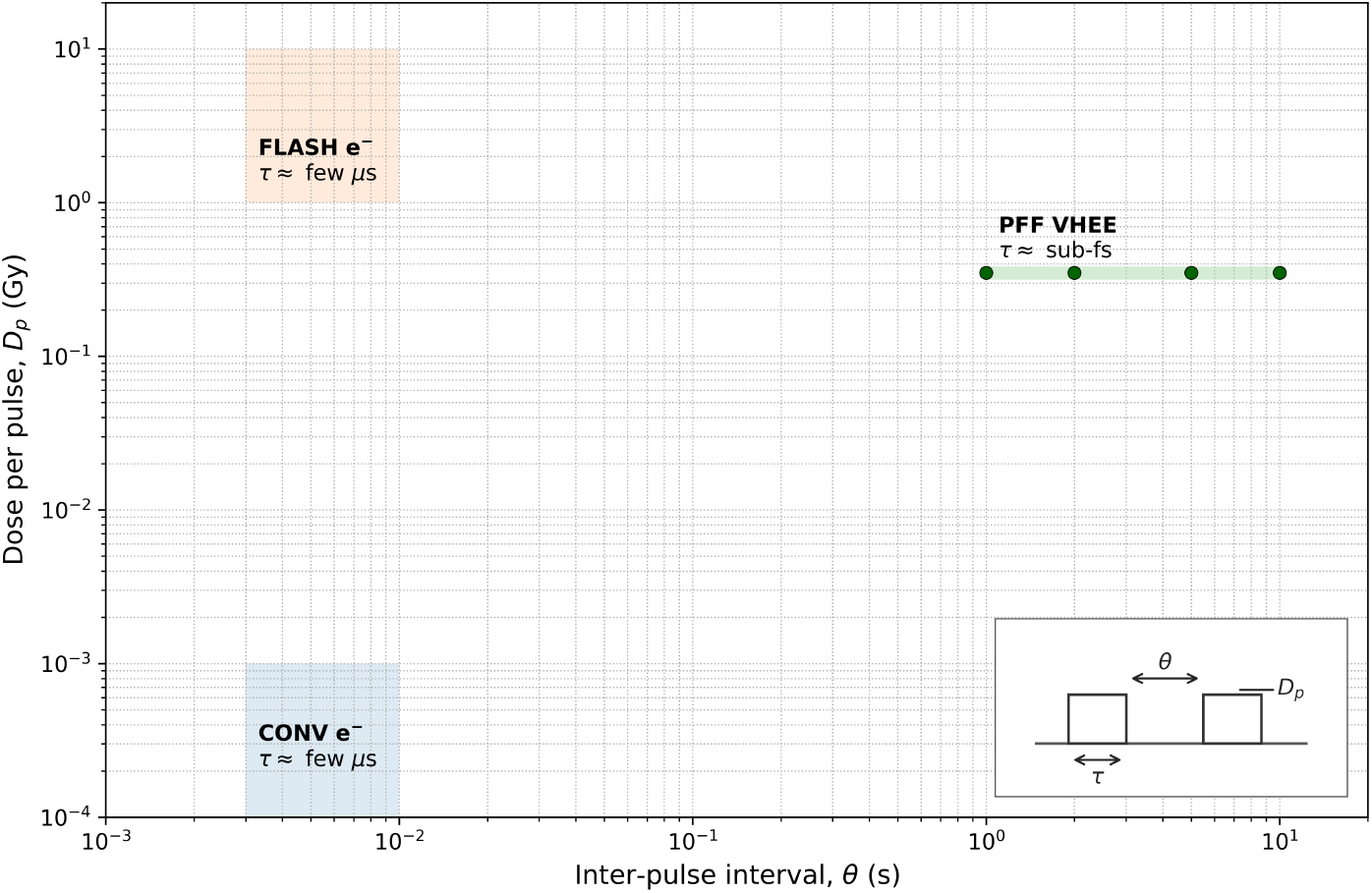
Conceptual map of electron-beam temporal microstructure in the (*θ, D*_*p*_) plane. Shaded regions schematically illustrate representative operating ranges for RF linac-based conventional and FLASH electron delivery. The PFF VHEE conditions investigated here occupy a distinct domain at second-scale inter-pulse intervals; markers indicate the values of *θ* explored in this work (*θ* = 1 s, 2 s, 5 s, 10 s). For each biological experiment, *θ* was varied while dose per pulse *D*_*p*_ and the prescribed total dose were held constant. Here, “pulse” denotes the biologically relevant dose-delivery event (a macropulse for RF linacs; one electron bunch for the LPA-driven VHEE source). The inset defines pulse duration *τ*, inter-pulse interval *θ*, and dose per pulse *D*_*p*_.

Here, we investigate the biological response to PFF using laser-driven VHEEs spanning 50 MeV to 300 MeV. Inter-pulse intervals *θ* = 1 s, 2 s, 5 s, and 10 s were investigated while maintaining the prescribed total dose and dose per pulse constant within each experimental system. Responses were evaluated in vitro in non-tumour human fibroblasts (MRC5-hTERT) and colorectal carcinoma cells (HCT116), and in vivo using zebrafish embryos. We investigated whether second-scale inter-pulse timing modulates biological response, whether this dependence differs between tumour and non-tumour cell models, and whether a timing-dependent response extends to an in vivo system.

## II. MATERIALS AND METHODS

### A. Laser-driven VHEE beam generation and delivery

VHEE irradiations were performed using a laser-driven electron source, as described in detail in Supplementary Section A1. In the present LPA configuration, each laser shot generated one electron bunch, corresponding to one irradiation pulse at the biological target. Here-after, “electron bunch” refers to the physical packet of accelerated electrons, whereas “pulse” refers to the corresponding dose-delivery event. The source generated sub-picosecond, broad-spectrum electron bunches with energies up to 300 MeV. Biological samples were positioned 16 cm downstream of the beam exit window in a dedicated holder, with fixed thickness diffusers used to improve lateral dose homogeneity across the irradiated area. Dose delivery was monitored in real time using a Razor Nano ionization chamber (IBA Dosimetry) positioned immediately downstream of the biological sample and independently verified using calibrated Gafchromic EBT-4 radiochromic films (RCF, Ashland, Bridgewater, NJ, USA) . The pulse repetition rate was varied from 1 Hz to 0.1 Hz, corresponding to inter-pulse intervals of *θ* = 1 s, 2 s, 5 s, and 10 s. Within each experiment, the dose per pulse was maintained constant across inter-pulse intervals, while the number of pulses was adjusted to deliver the prescribed total dose. Electron spectra were acquired immediately before and after each irradiation condition to monitor accelerator stability and assess potential spectral drift. The measured spectra were subsequently combined with Monte Carlo-derived energy-dependent dose-deposition profiles to characterise the dose-weighted electron energy distribution at the biological target. Representative dose-weighted spectra are shown in Supplementary Figure A2.

The resulting dose-weighted mean electron energy was approximately 136 MeV and remained stable across irradiation conditions and experimental days, with variations not exceeding 7 %. For a 350 mGy pulse, 90 % of the dose within the biological region of interest was deposited within approximately 150 fs, corresponding to an effective instantaneous dose rate on the order of 10^12^ Gy s^*−*1^ (Supplementary Figure A3). Further details of the experimental setup and irradiation geometry are provided in Supplementary Section A1, while dosimetric, spectral, and temporal dose-delivery characterisation are reported in Supplementary Section A2.

### B. In vitro cell irradiation and viability assessment

The immortalized human fibroblast cell line MRC5-hTERT and the human colorectal carcinoma cell line HCT116 were used for the in vitro experiments. MRC5-hTERT cells were cultured in Dulbecco’s Modified Eagle Medium (DMEM + GlutaMAX, Gibco), and HCT116 cells in McCoy’s 5A (Modified) Medium (Gibco). Both media were supplemented with 10 % fetal calf serum (PAA) and 1 % Penicillin–Streptomycin (Thermo Fisher Scientific). Cells were maintained as monolayers at 37 °C in a humidified atmosphere containing 5 % CO_2_, following the protocol described by Bayart et al.[9].

For irradiation, 10^4^ cells were transferred into 2 mL Eppendorf tubes containing growth medium, centrifuged, and irradiated at room temperature (22 °C – 24 °C) to a total dose of 4.5 Gy. The four inter-pulse interval conditions were applied while maintaining the total dose and dose per pulse constant, with mean doses per pulse of 0.36 *±* 0.06 Gy for MRC5-hTERT and 0.32 *±* 0.03 Gy for HCT116. All four conditions were tested independently on three separate experimental days.

Following irradiation, each pellet was re-suspended and seeded evenly into four wells of a 12-well plate, with fresh medium added to a final volume of 2 mL per well. Cells were incubated for five generations (six days for MRC5-hTERT; five days for HCT116), then harvested using 0.2 mL Accutase (EMD Millipore) and neutralized with an equal volume of growth medium. Viable cells in each well were counted using an ORFLO Moxi Mini Automated Cell Counter (Type S cassette), and the mean of the four wells was used as the viability endpoint for each independent experiment.

### C. In vivo zebrafish irradiation and morphological assessment

Wild-type AB zebrafish embryos were maintained in E3 medium at 28 °C until irradiation. At a developmental stage corresponding to 4 h post-fertilization (hpf), embryos were transferred into 2 mL Eppendorf tubes containing E3 medium and irradiated at room temperature (22 °C – 24 °C) to a total dose of 10 Gy. The four inter-pulse interval conditions were applied while maintaining the total dose and dose per pulse constant, with a mean dose per pulse of 0.35*±*0.04 Gy. All four conditions were tested independently on three separate experimental days, with an average of 56 embryos per condition. At 5 days post-irradiation (dpi), when major organogenesis is complete [10] and embryos are not yet considered protected animals under European Directive 2010/63/EU [11], embryos were fixed in 10 % formalin and imaged using an Echo Rebel microscope equipped with a 4*×* objective. Morphological assessment was based on body length as an indicator of growth impairment, quantified using a custom Python-based image-analysis tool along the body axis from the anterior tip of the head to the posterior end of the vertebral column, excluding the caudal fin fold.

### D. Statistical analysis

For both cell viability and zebrafish assays, data were normalized to the corresponding non-irradiated (NI) controls from the same experimental day. Statistical comparisons across inter-pulse interval conditions and, where applicable, between biological models were performed using the Kruskal–Wallis test, followed by Dunn’s post hoc test with Benjamini– Hochberg false discovery rate (FDR) correction. All tests were two-sided, and *p*-value *<* 0.05 was considered statistically significant. Reported *p*-values for post hoc comparisons are FDR-adjusted. Data are presented as mean *±* standard error of the mean (SEM), unless otherwise stated. All analyses were performed using GraphPad Prism (version 10.3.1).

## III. RESULTS

### A. In vitro cell viability under temporally modulated VHEE irradiation

MRC5-hTERT fibroblasts and HCT116 colorectal carcinoma cells were irradiated to a total dose of 4.5 *±* 0.2 Gy across irradiation conditions. Cell viability showed a distinct dependence on inter-pulse interval in the two cell lines (Figure 2). In MRC5-hTERT fibroblasts, viability was highest at the shortest inter-pulse interval (*θ* = 1 s) and was significantly higher than at 2 s (*p <* 0.001), 5 s (*p <* 0.01), and 10 s (*p <* 0.01). In contrast, HCT116 cells showed the highest viability at the longest inter-pulse interval (*θ* = 10 s), which was significantly higher than at all shorter intervals (1 s, 2 s, and 5 s, all *p <* 0.05). No significant differences were observed among the remaining intervals within either cell line. Direct comparison between the two cell lines showed significantly higher viability in MRC5-hTERT fibroblasts than in HCT116 cells at *θ* = 1 s (*p <* 0.01). Overall, these results show distinct inter-pulse interval response patterns between the two cell lines, with maximal viability occurring at opposite ends of the investigated temporal range. Exact FDR-adjusted *p*-values for within-cell-line comparisons are reported in Supplementary Tables A1 and A2, while between-cell-line comparisons are reported in Supplementary Table A3.

**FIG. 2.**
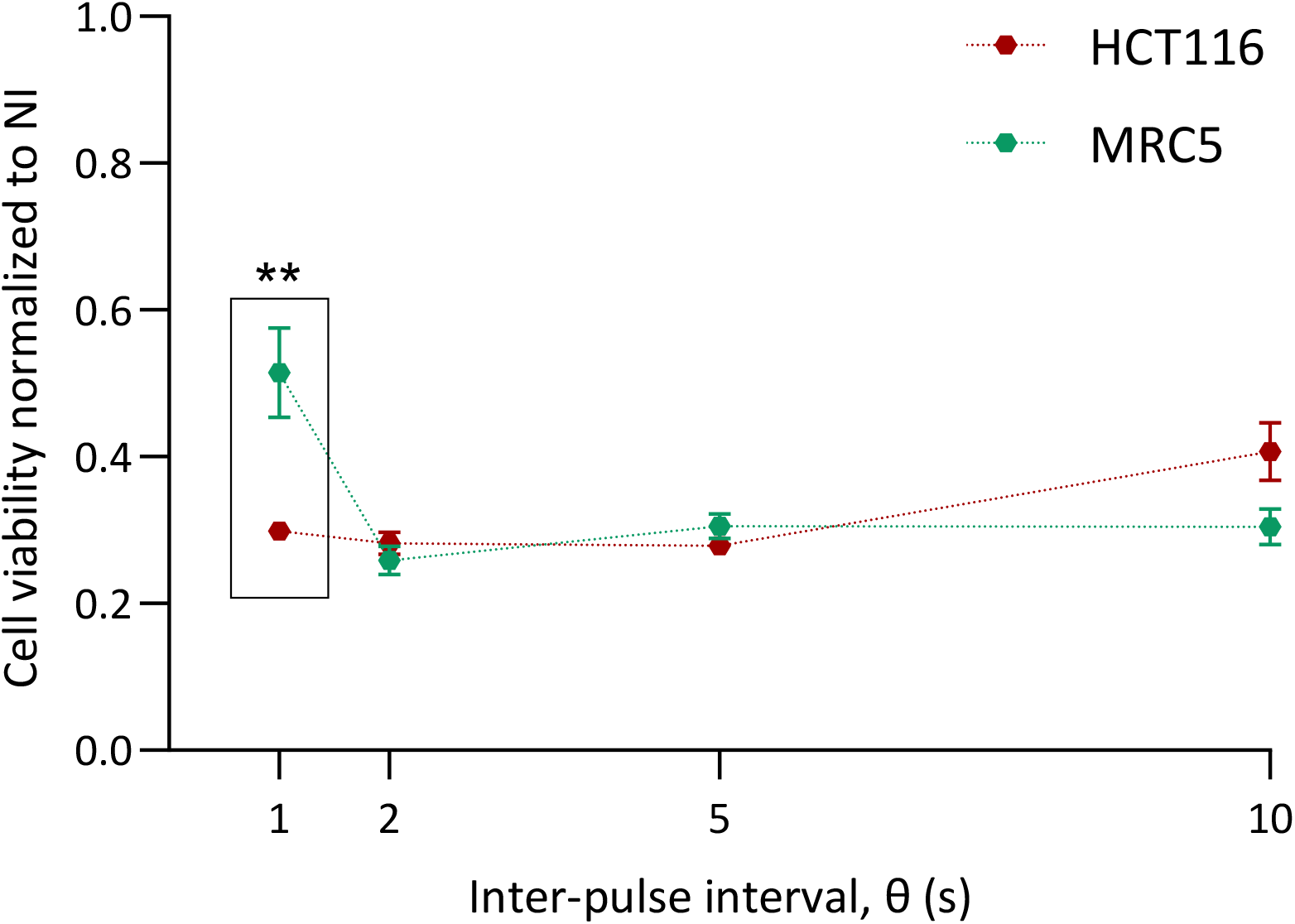
Normalized cell viability as a function of inter-pulse interval (*θ*) for MRC5-hTERT fibroblasts and HCT116 colorectal carcinoma cells following laser-driven VHEE irradiation (total dose 4.5 *±* 0.2 Gy). Each point represents the mean normalized cell viability ± SEM from three independent experiments. The boxed region highlights the between-cell-line comparison at *θ* = 1 s. Statistical testing: Kruskal-Wallis followed by Dunn’s post hoc test with Benjamini-Hochberg FDR correction. Significance levels: ns, not significant; ∗ p *<* 0.05; ∗∗ p *<* 0.01; ∗∗∗ p *<* 0.001; ∗∗∗∗ p *<* 0.0001.

### B. In vivo developmental response under temporally modulated VHEE irradiation

Wild-type AB zebrafish embryos were irradiated to a total dose of 10.2 *±* 0.4 Gy across irradiation conditions. Normalized body length at 5 dpi showed a pronounced dependence on inter-pulse interval (Figure 3). Embryos irradiated at the shortest inter-pulse interval (*θ* = 1 s) exhibited significantly greater body length than those irradiated at 2 s, 5 s, and 10 s (all *p <* 0.0001), indicating reduced growth impairment at the shortest inter-pulse interval. A smaller but significant difference was also observed between the 5 s and 10 s conditions (*p* = 0.0438); the remaining comparisons among the longer intervals were not significant. Exact FDR-adjusted *p*-values for pairwise comparisons are reported in Supplementary Table A4. Notably, this response pattern was consistent with that observed in MRC5-hTERT fibroblasts, with both models showing their highest respective endpoint at *θ* = 1 s.

**FIG. 3.**
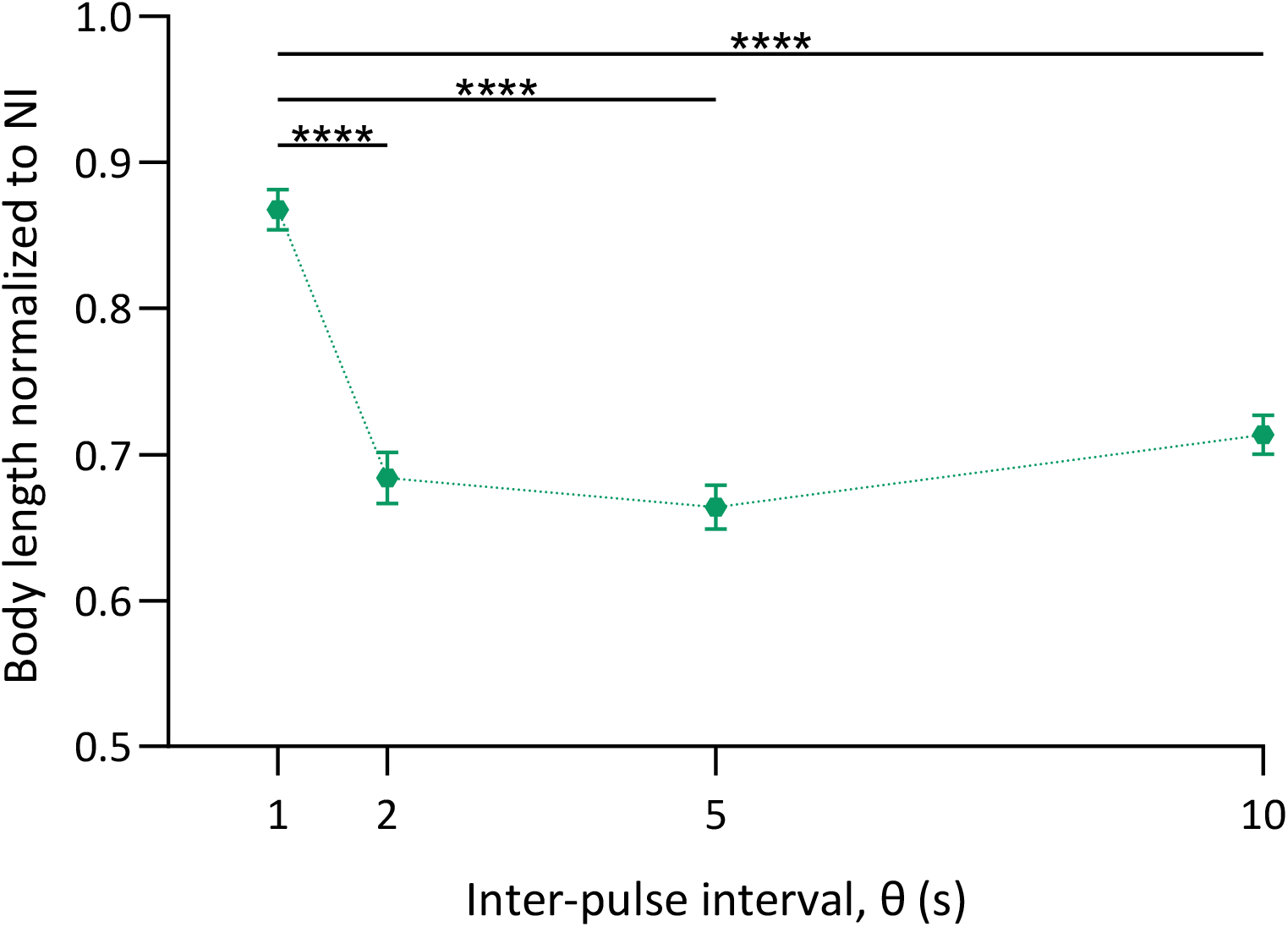
Normalized zebrafish body length at 5 dpi as a function of the inter-pulse interval (*θ*), following laser-driven VHEE irradiation (total dose 10.2 *±* 0.4 Gy). Body length was normalized to the corresponding NI control from the same experimental day. Data are presented as mean *±* SEM across three independent experiments. Statistical testing: Kruskal–Wallis test with Dunn’s post hoc with Benjamini-Hochberg FDR correction. Significance levels: ns, not significant; ∗ p *<* 0.05; ∗∗ p *<* 0.01; ∗∗∗ p *<* 0.001; ∗∗∗∗ p *<* 0.0001.

## IV. DISCUSSION

This study shows that second-scale inter-pulse timing can influence the biological response to laser-driven VHEE irradiation when total dose and dose per pulse are maintained constant across irradiation conditions. MRC5-hTERT fibroblasts and HCT116 colorectal carcinoma cells exhibited distinct temporal responses, with the shortest investigated interval associated with higher viability in MRC5-hTERT fibroblasts but lower viability in HCT116 cells. Notably, this same interval was associated with reduced radiation-induced growth impairment in zebrafish embryos, extending the timing-dependent response observed in the non-tumour cell model to an in vivo system. Together, these findings identify inter-pulse timing as a biologically relevant parameter of pulsed VHEE irradiation and highlight its potential to differentially modulate responses in non-tumour and tumour systems.

Evidence from FLASH irradiation has highlighted the importance of the temporal characteristics of dose delivery in radiobiological response. Yet, temporal delivery regimes combining ultrashort dose-deposition events with overall irradiation times extending over seconds remain comparatively underexplored. FLASH sparing has been reported across different radiation qualities [12–15], but the relative contributions of dose per pulse, instantaneous and average dose rate, and overall irradiation time remain unresolved [16, 17]. In particular, it remains unclear to what extent the biological response is determined by individual UHDR dose-deposition events or by the temporal characteristics of the complete irradiation. The PFF conditions investigated here provide access to this distinct combination of temporal scales: individual VHEE irradiation pulses deposited their dose within the sub-picosecond range, corresponding to effective instantaneous dose rates on the order of 10^12^ Gy s^*−*1^, while successive pulses were separated by 1 s to 10 s. Thus, despite the ultra-high instantaneous dose rate within each pulse, the complete irradiation extended over seconds and the corresponding average dose rate remained below the range typically associated with FLASH delivery (tens of Gy s^*−*1^).

The temporal response observed here is consistent with previous evidence that radiation response can depend on pulse timing. In our previous work with laser-driven protons, varying the interval between ultrashort radiation pulses over a broader but overlapping second-scale range produced a non-monotonic survival response in cancer cell models [9]. Notably, HCT116 cells were investigated in both studies and showed a comparable dependence on inter-pulse timing, with higher survival observed at longer intervals following both laser-driven proton and VHEE irradiation within the overlapping time window. The recurrence of this temporal response in the same cell line with two different particle beams indicates that the phenomenon is not specific to VHEE and provides evidence that temporal sensitivity can occur across distinct radiation qualities. Earlier high-dose-rate photon and electron studies also reported non-monotonic survival responses [18, 19], further supporting the broader relevance of temporal structure in radiobiological response.

The biological mechanisms underlying this temporal dependence remain to be established. Mechanistic findings from our previous proton PFF study in HCT116 cells may nevertheless provide relevant insight: the non-monotonic response to inter-pulse timing was dependent on PARP1 activity [9], supporting a role for DNA damage processing in the temporal response. This finding echoes earlier observations of the W-effect, in which varying the interval between two UHDR radiation pulses produced a non-monotonic variation in radiosensitivity [20, 21]. However, the irradiation schemes differ substantially. Classical W-effect experiments involved two radiation pulses, whereas PFF in the present study comprised approximately a dozen to several tens of pulses, depending on the biological model. This difference in pulse number may contribute to the distinct temporal response observed under PFF compared with that reported for the W-effect. While these observations support DNA damage processing as a candidate mechanism, other transient processes, including oxygen chemistry and redox regulation, may also contribute to the dependence on inter-pulse timing. Differences in these processes between non-tumour and tumour cells could, in principle, contribute to the contrasting temporal responses observed in MRC5-hTERT and HCT116 cells. Dedicated mechanistic studies will be required to determine their respective contributions. Varying the inter-pulse interval necessarily altered both the overall irradiation time and the average dose rate, while total dose, dose per pulse, and instantaneous dose rate were maintained constant across irradiation conditions. Average dose rate therefore remains intrinsically coupled to inter-pulse timing and may contribute to the observed biological response. Previous observations in MRC5-hTERT cells nevertheless provide relevant context for the response observed at the shortest intervals. In our previous study [8], comparable radiosensitivity was observed following pulsed laser-driven VHEE irradiation at approximately 12 Gy*/*min and conventional 7 MeV linac-generated electron irradiation at approximately 30 Gy*/*min. In the present study, the *θ* = 1 s and *θ* = 2 s conditions corresponded to average dose rates of approximately 20 Gy*/*min and 10 Gy*/*min, respectively, yet produced significantly different responses in MRC5-hTERT cells. These observations suggest that average dose rate alone does not provide a straightforward explanation for the MRC5-hTERT response at short inter-pulse intervals. The absence of a corresponding difference between these conditions in HCT116 cells further emphasizes the dependence of the temporal response on the biological model. Nevertheless, because inter-pulse interval, average dose rate, and overall irradiation time remain coupled, their respective contributions cannot be fully disentangled from the present experiments.

Beyond the contrasting responses observed in the tumour and non-tumour cell models, the zebrafish experiments show that the biological response to inter-pulse timing extends to an in vivo vertebrate model. The greater body length observed at *θ* = 1 s indicates reduced radiation-induced growth impairment at the same inter-pulse interval associated with the highest viability in MRC5-hTERT fibroblasts. Thus, despite the different biological end-points, both non-tumour models showed the lowest radiation-induced effect at the shortest investigated interval. This consistency across an in vitro cellular model and an in vivo vertebrate model provides evidence that the biological influence of second-scale pulse timing extends beyond the cellular level. Further studies will be required to establish the generality of these observations across additional normal and tumour models spanning different levels of biological complexity, clinically relevant dose levels, complementary biological endpoints, and longer-term functional outcomes. Alternative pulse-delivery schemes will be required to disentangle the respective contributions of inter-pulse timing, average dose rate, and overall irradiation time, which remain coupled under the conditions investigated here. Direct comparisons with FLASH-VHEE delivery will further help establish how different temporal characteristics of dose delivery contribute to biological response and whether they can be exploited to enhance differential responses between normal and tumour systems.

## V. CONCLUSIONS

This study demonstrates that biological response to laser-driven VHEE irradiation can be modulated by the second-scale temporal separation of ultrashort, ultra-high instantaneous dose-rate pulses, while maintaining total dose and dose per pulse constant. Distinct temporal responses were observed in non-tumour MRC5-hTERT fibroblasts and HCT116 colorectal carcinoma cells, while the response observed in zebrafish embryos provides in vivo evidence that sensitivity to inter-pulse timing extends beyond isolated cell systems. Notably, the shortest investigated interval (*θ* = 1 s) was associated with reduced radiation-induced effects in both MRC5-hTERT fibroblasts and zebrafish embryos, while a contrasting response was observed in HCT116 cells.

Together, these findings identify inter-pulse timing as a biologically relevant parameter of pulsed VHEE delivery and establish PFF as a distinct temporal irradiation regime enabled by laser-plasma accelerators. Although the respective contributions of inter-pulse timing, average dose rate, and overall irradiation time remain to be disentangled, the differential responses observed across biological models motivate further investigation of temporal pulse structure as a potential means of modulating radiobiological response.

