## Supplementary Material for "Laser-driven VHEE pulsed fast fractionation (PFF): second-scale inter-pulse timing can modulate normal tissue toxicity in vivo"

### A1. LASER-DRIVEN VHEE SOURCE AND IRRADIATION SETUP

VHEE beams were generated by laser wakefield acceleration (LWFA) using a 60 TW Ti:sapphire laser system delivering 2.4 J, 30 fs full width at half maximum (FWHM) pulses on target. The laser was focused by an $f/18$ off-axis parabolic mirror onto a helium–nitrogen gas jet containing 2 % $N_2$, delivered through a 4 mm × 250 µm rectangular nozzle (Figure A1). The laser–plasma interaction generated broadband electron bunches with energies extending up to approximately 300 MeV and a duration of approximately 10 fs. The total charge in each electron bunch (irradiation pulse) exceeded 500 pC and was monitored using a fast current transformer (Turbo-ICT, Bergoz, Saint-Genis-Pouilly, France).

The experimental setup was operated in two configurations (Figure A1). In diagnostic mode, electron spectra were measured using a magnetic spectrometer comprising a 1 T dipole magnet and a LANEX scintillating screen imaged by a CCD camera. In irradiation mode, the dipole magnet was removed from the beam path, allowing the electron beam to exit the vacuum chamber through a 1.5 mm aluminium window and propagate 16 cm in air to the irradiation plane. Biological samples were positioned in a dedicated polymethyl methacrylate (PMMA) holder accommodating a 2 mL tube.

Electron spectra were acquired immediately before (PRE) and after (POST) each biological irradiation for all investigated inter-pulse conditions and experimental days. Each PRE and POST measurement comprised 30 consecutive electron bunches. These measurements were used to monitor the stability of the laser-plasma accelerator and to verify that no substantial spectral drift occurred during the irradiation campaign. The corresponding dose-weighted spectral analysis is described in Section A2.2.

### A2. DOSIMETRIC CHARACTERISATION OF THE VHEE BEAM

#### A2.1. Dose monitoring and spatial dose distribution

During VHEE irradiations, delivered dose was monitored in real time using a Razor Nano ionisation chamber (IC; IBA Dosimetry, Schwarzenbruck, Germany) positioned immediately

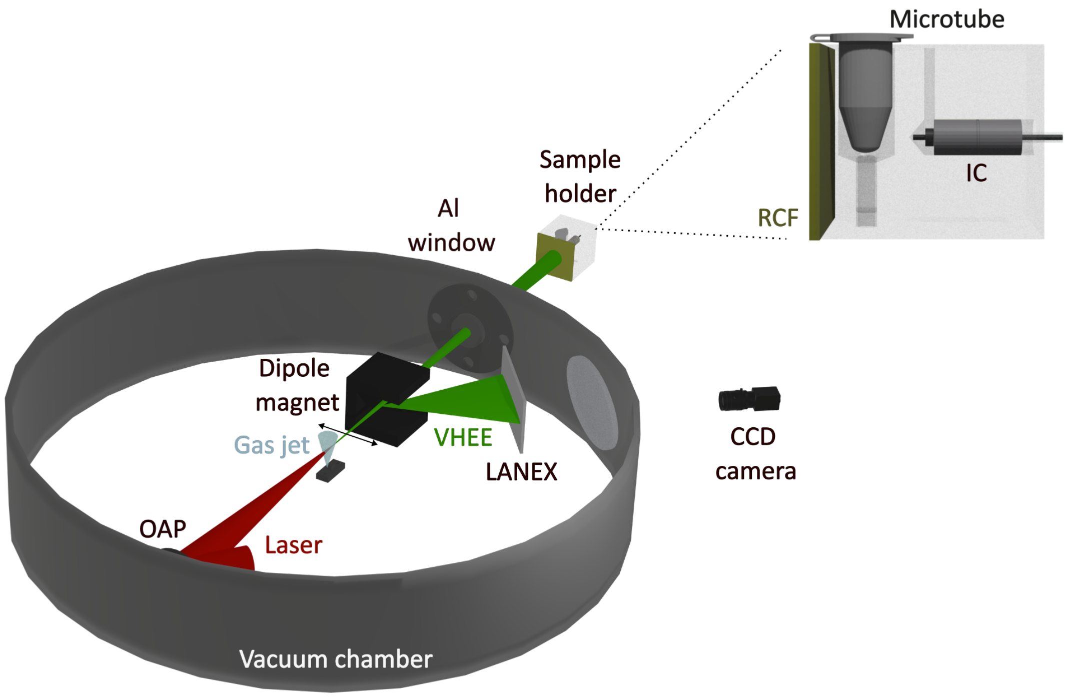


FIG. A1. Top view of the LPA experimental setup. The laser is focused by an off-axis parabolic mirror (OAP) onto a He/$N_2$ gas-jet target, generating the VHEE electron beam. In diagnostic mode, a dipole magnet disperses the electron beam onto a LANEX scintillating screen for spectral characterization. In irradiation mode, the dipole is removed and the beam exits the vacuum chamber through a 1.5 mm Al window before reaching the biological sample holder. The enlarged view shows the arrangement of the radiochromic film (RCF), sample microtube, and ionisation chamber (IC) within the sample holder.

downstream of the biological sample (Figure A1). Prior to each experimental session, a calibration was performed to establish the relationship between the dose measured at the IC position and that delivered at the sample plane. This calibration was used to determine the dose per pulse ($D_p$) at the sample plane and the number of electron pulses required to reach the prescribed total dose. Beam divergence was increased using a 1 mm brass diffuser mounted on the exit window. At the biological target plane, the resulting beam divergence was $61.2 \pm 2.2$ mrad for the in vitro irradiation configuration and $81.0 \pm 1.6$ mrad for the in vivo zebrafish configuration. Biological samples were positioned behind a fixed 1 cm water-equivalent absorber. Together, these passive elements provided a broader lateral dose distribution and improved dose homogeneity across the sample plane.

Gafchromic EBT-4 radiochromic films (RCF; Ashland, Bridgewater, NJ, USA) were positioned in front of the biological samples to independently verify the delivered dose and characterise its spatial distribution across the irradiation plane. Prior to the irradiation experiments, the RCFs were absolutely calibrated using a 7 MeV clinical electron beam at Institut Curie (Paris, France), following the protocol described in [1]. RCF measurements were also used to estimate the uncertainty associated with the delivered dose, which was propagated across experimental replicates in the reported dose values. The dominant contribution arose from transverse dose non-uniformity across the irradiated area, whereas longitudinal, depth-dependent variations were negligible under the employed irradiation conditions.

#### A2.2. Dose-weighted VHEE spectral characterisation

To assess the effective energy distribution of the broadband VHEE beam at the biological target, the measured electron spectra were combined with Monte Carlo-derived energy-dependent dose deposition within a circular 5 mm-diameter region of interest (ROI) corresponding to the transverse extent of the biological target. Each spectral component was weighted according to its relative contribution to the dose deposited within this ROI, and a dose-weighted mean electron energy, $E_w$, was calculated from the resulting distributions. This analysis provided the dose-weighted electron energy distribution at the biological target, confirming the dominant contribution of electrons in the VHEE energy range. Representative dose-weighted spectra and the corresponding $E_w$ values for the four inter-pulse intervals during one experimental day are shown in Figure A2. For each condition, the displayed distribution corresponds to the mean of the PRE- and POST-irradiation dose-weighted spectra.

Across all irradiation conditions and experimental days, variations in $E_w$ between PRE and POST measurements remained below 6 %. Within individual experimental days, the maximum variation in $E_w$ among inter-pulse conditions, evaluated from PRE/POST-averaged spectra, was below 7 %. Comparison of measurements acquired at the same inter-pulse interval across independent experimental days showed a relative dispersion below 5 %. These measurements indicate that the dose-weighted spectral characteristics at the biological target remained stable throughout the irradiation campaign.

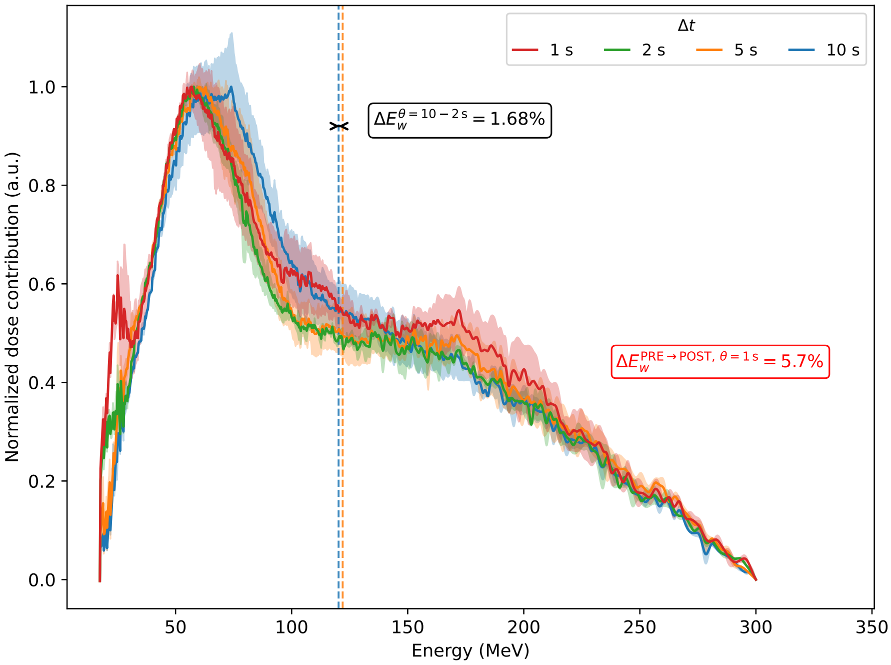


FIG. A2. Dose-weighted electron energy spectra for the four investigated inter-pulse intervals ($\theta = 1\,\mathrm{s}$, $2\,\mathrm{s}$, $5\,\mathrm{s}$, and $10\,\mathrm{s}$) during a representative experimental day. For each condition, the solid line represents the mean of the PRE- and POST-irradiation dose-weighted spectra, normalized to its maximum, while the shaded region spans the PRE–POST range. Vertical dashed lines indicate the corresponding dose-weighted mean energies, $E_w$. Annotations indicate the maximum PRE–POST relative change and the maximum inter-condition variation in $E_w$.

#### A2.3. Temporal dose-deposition characteristics

The temporal profile of dose deposition at the biological target was calculated using time-of-flight (ToF) calculations applied to the experimental electron spectra. The ToF calculation accounts for the energy-dependent arrival times of the different spectral components at the target plane, providing an estimate of the temporal distribution of electrons reaching the biological target. To derive the corresponding temporal distribution of dose deposition, each spectral component was additionally weighted according to its energy-dependent contribution to the dose deposited within the same 5 mm-diameter ROI used for the dose-weighted spectral analysis in Section A2.2. As shown in Figure A3, the resulting dose-deposition profile is concentrated at earlier times than the overall electron-arrival distribution, indicating that the earlier-arriving spectral components contribute preferentially to dose deposition within the biological target. The time coordinate was defined relative to the arrival of the first electrons at the target plane. The dose-deposition time, $t_{90}$, was defined as the time required for the cumulative dose-deposition distribution to reach $90\,\%$ of the total deposited dose. The resulting $t_{90}$ was approximately $150\,\mathrm{fs}$. For a representative dose per pulse of $D_p = 350\,\mathrm{mGy}$, this corresponds to an effective instantaneous dose rate of approximately $2.1 \times 10^{12}\,\mathrm{Gy\,s^{-1}}$.

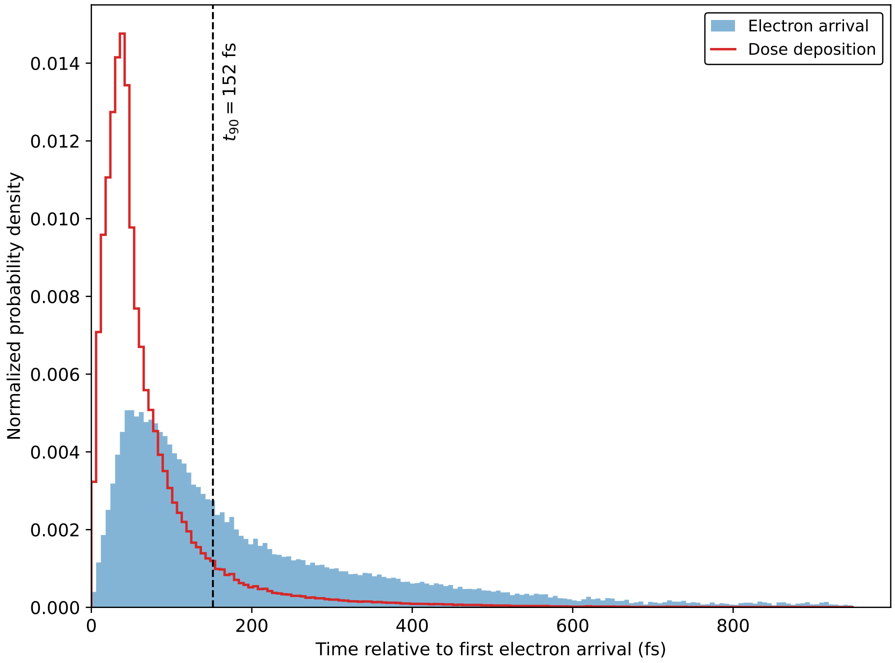


FIG. A3. Temporal distributions of electron arrival and dose deposition within the $5\,\mathrm{mm}$-diameter ROI corresponding to the biological target. Both distributions are normalized to unit integral, and time is expressed relative to the arrival of the first electrons. The dashed line indicates the calculated $t_{90} = 152\,\mathrm{fs}$.

### A3. SUPPLEMENTARY STATISTICAL ANALYSES

#### A3.1. In vitro cell viability

Within each cell line, pairwise comparisons among the four investigated inter-pulse intervals were performed separately for MRC5-hTERT and HCT116 cells using Dunn's post hoc test following the Kruskal–Wallis test, with Benjamini–Hochberg false discovery rate (FDR) correction. Exact FDR-adjusted $p$-values are reported in Tables A1 and A2, respectively.

To further examine the contrasting responses observed at the extremes of the investigated temporal range, an additional analysis was performed including the 1 s and 10 s conditions from both cell lines. Pairwise comparisons from this analysis were performed using Dunn's post hoc test with Benjamini–Hochberg FDR correction. Between-cell-line comparisons are reported in Table A3.

TABLE A1. Pairwise comparisons of normalized cell viability among inter-pulse intervals for MRC5-hTERT fibroblasts.

| Comparison | Adjusted $p$-value | Significance |
|---|---|---|
| 1 s vs 2 s | 0.0002 | *** |
| 1 s vs 5 s | 0.0082 | ** |
| 1 s vs 10 s | 0.0082 | ** |
| 2 s vs 5 s | 0.2651 | ns |
| 2 s vs 10 s | 0.2651 | ns |
| 5 s vs 10 s | 0.9238 | ns |

ns, not significant; *$p < 0.05$; **$p < 0.01$; ***$p < 0.001$.

TABLE A2. Pairwise comparisons of normalized cell viability among inter-pulse intervals for HCT116 colorectal carcinoma cells.

| Comparison | Adjusted $p$-value | Significance |
|---|---|---|
| 1 s vs 2 s | 0.6425 | ns |
| 1 s vs 5 s | 0.6299 | ns |
| 1 s vs 10 s | 0.0497 | * |
| 2 s vs 5 s | 0.8518 | ns |
| 2 s vs 10 s | 0.0132 | * |
| 5 s vs 10 s | 0.0132 | * |

ns, not significant; *$p < 0.05$; **$p < 0.01$; ***$p < 0.001$.

TABLE A3. Between-cell-line comparisons of normalized cell viability at the two extremes of the investigated inter-pulse interval range ($\theta = 1\,\mathrm{s}$ and $10\,\mathrm{s}$).

| MRC5-hTERT | HCT116 | Adjusted $p$-value | Significance |
|---|---|---|---|
| 1 s | 1 s | 0.0023 | ** |
| 1 s | 10 s | 0.2875 | ns |
| 10 s | 1 s | 0.6084 | ns |
| 10 s | 10 s | 0.1097 | ns |

ns, not significant; *$p < 0.05$; **$p < 0.01$; ***$p < 0.001$; ****$p < 0.0001$.

#### A3.2. In vivo zebrafish response

Pairwise comparisons of normalized body length among inter-pulse intervals were performed using Dunn's post hoc test following the Kruskal–Wallis test, with Benjamini–Hochberg FDR correction. Exact FDR-adjusted $p$-values are reported in Table A4.

TABLE A4. Pairwise comparisons of normalized body length among inter-pulse intervals in irradiated zebrafish embryos.

| Comparison | Adjusted $p$-value | Significance |
|---|---|---|
| 1 s vs 2 s | $< 0.0001$ | **** |
| 1 s vs 5 s | $< 0.0001$ | **** |
| 1 s vs 10 s | $< 0.0001$ | **** |
| 2 s vs 5 s | 0.3849 | ns |
| 2 s vs 10 s | 0.3169 | ns |
| 5 s vs 10 s | 0.0438 | * |

ns, not significant; *$p < 0.05$; **$p < 0.01$; ***$p < 0.001$; ****$p < 0.0001$.

[1] L. Giuliano, G. Franciosini, L. Palumbo, L. Aggar, M. Dutreix, L. Faillace, V. Favaudon, G. Felici, F. Galante, A. Mostacci, M. Migliorati, M. Pacitti, A. Patriarca, and S. Heinrich, Characterization of Ultra-High-Dose Rate Electron Beams with ElectronFlash Linac, Applied Sciences **13**, 631 (2023).
